# Impact of Pericytes on the Stabilisation of Microvascular Networks in Microfluidic Systems in Response to Nanotoxicity

**DOI:** 10.1101/2022.05.03.490457

**Authors:** Matthew Dibble, Stefania Di Cio, Piaopiao Luo, Frances Balkwill, Julien E. Gautrot

**Affiliations:** Institute of Bioengineering, University of London, Mile End Road, London, E1 4NS, UK; School of Engineering and Materials Science, Queen Mary, University of London, Mile End Road, London, E1 4NS, UK; Barts Cancer Institute, Queen Mary University of London, Charterhouse Square, London EC1M 6BQ

## Abstract

Recapitulating the normal physiology of the microvasculature is pivotal in the development of more complex in vitro models and organ-on-chip design. Pericytes are an important component of the vasculature, promoting vessel stability, inhibiting vascular permeability and maintaining the vascular hierarchical architecture. This report presents a microfluidic model exploring interactions between endothelial cells and pericytes. We identify basal conditions required to form stable and reproducible endothelial networks. We then investigate interactions between endothelial cells and pericytes via direct co-culture. In our system, pericytes inhibited vessel hyperplasia and maintained vessel length in prolonged culture (>10 days). In addition, these vessels displayed barrier function and expression of junction markers associated with vessel maturation, including VE-cadherin, β-catenin and ZO-1. Furthermore, pericytes maintained vessel integrity following stress (nutrient starvation) and inhibited vessel regression, in contrast to the striking dissociation of networks in endothelial monocultures. This response was also observed when endothelial/pericyte co-cultures were exposed to high concentrations of moderately toxic cationic nanoparticles used for gene delivery. This study highlights the importance of pericytes in protecting vascular networks from stress and external agents and their importance to the design of advanced *in vitro* models, including for the testing of nanotoxicity, to better recapitulate physiological response and avoid false positives.

## 1. Introduction

The vasculature is an integral component of many physiological and pathophysiological conditions and its formation and stabilisation *in vitro* is essential for the development of vascularised tissue models [1-5]. Such 3D *in vitro* models are particularly important for the accurate prediction of drug and nanomaterial toxicity, particularly considering the ubiquity of particle exposure (metallic, ceramic and polymeric) in daily life [6, 7]. Traditional 2D systems lack physiological characteristics (geometry, mechanical and biochemical stimuli) which might lead to misrepresentation of normal physiological *in vivo* processes and their response to fine chemicals, therapeutic agents and nanomaterials. Although 2D models have been useful in identifying the potential impact of such compounds on endothelial biology and barrier function of the endothelium [8], and remain more adapted to high throughput screening, 3D models emerge as particularly useful alternatives for more accurate prediction of safety and efficacy testing [2, 9]. The emergence of organ-on-a-chip systems is crucial in this effort and allows, for example, the addition of flow and biochemical gradients [10]. Atmospheric nanoparticles were found to increase vessel permeability by disrupting barrier functions in a 3D vasculature-on-a-chip model [11]. Using another 3D microvessel-on-a-chip system, the extravascular transport mechanisms of cationic polymer nanoparticles was assessed [12].

Researchers have developed various iterations of a ‘vasculature-on-a-chip’ [3, 13-20]. These models generally use PDMS chips which are manufactured using photo- and soft-lithography. A relatively established design features a central gel compartment flanked by lateral medium compartments, separated by microposts. The central gel compartment is injected with a hydrogel precursor, together with endothelial cells, following by culture to allow the assembly of a microvasculature and the maturation of its lumenised structure. Microfluidic devices offer several advantages over more traditional *in vitro* blood vessel models, including co-culture potential, real-time imaging, and physiological architecture. However, some of the parameters regulating the formation of such microvasculature on chips (concentrations of fibrinogen, thrombin, aprotinin, endothelial cell density, presence of stromal cells and exogenous factors) have not been studied systematically to rigorously compare and present their impact on network maturation [13, 16, 20-24]. Such variation in conditions can lead to poor reproducibility and makes direct comparison between different studies limited. A side-by-side comparison of the importance of these different factors is therefore important for the wider implementation of these models by the bioengineering community.

One common problem encountered when culturing microvasculatures for prolonged times (> 4 days) is vessel hyperplasia. This could limit long-term culture and the use of these systems for embedding spheroids and organoids for the development of advanced tissue models. To overcome this issue, endothelial cells can be cultured with pericytes [15-17]. Pericytes are abluminally located mesenchymal cells that have the capacity to differentiate in many types of cells [25] Due to their heterogeneity, the identification of specific markers has been disputed [26], but platelet-derived growth factor receptor-β (PDGFRβ) and neural/glial antigen-2 (NG2) are generally accepted markers associated with these cells [27, 28] Pericytes typically interact with a number of endothelial cells, with primary cytoplasmic processes running along the abluminal surface of the endothelium and secondary processes running perpendicularly, enclosing the endothelial tube [29]. Co-culturing endothelial cells with pericytes inhibits vessel hyperplasia and endothelial cell proliferation, and promotes vessel barrier function and endothelial cell survival [15, 17, 30-32].

In this study, we first examine the role of a range of parameters regulating the establishment of a stable microvasculature, including the effect of cell density, fibrinogen, collagen I, VEGF and aprotinin concentrations. In addition, we study how pericytes stabilise endothelial network formation and impact on its barrier function. We then apply this system to the study of toxicity to the microvasculature, in the context of nanoparticle response.

## 2. Materials and Methods

### 2.1. Microfabrication and material characterisation

#### Microfluidic chip fabrication

Microfluidic devices were fabricated using photo- and soft lithography. A master with positive relief patterns of SU-8 2050 photoresist (A-Gas Electronic Materials) on a silicon wafer (PI-KEM) was prepared by photolithography. A PDMS (Ellsworth Adhesives) polymer was cast against this master and thermally cured to obtain a negative replica piece. After separating from the master, hydrogel ports and medium reservoirs were punched from the PDMS stamp using biopsy punches. The PDMS stamp is then bonded to a glass coverslip using an oxygen plasma treatment. Devices were then autoclaved and dried at >60 °C for 3 days to restore hydrophobicity.

#### Contact angle goniometry

Contact angle goniometry was used to investigate PDMS hydrophobic recovery post-plasma treatment. Samples were exposed to a 5 µL deionised water droplet and the contact angle between the PDMS and water droplet was extrapolated using the ‘Default Method’ of the DSA 100 (Kruss Scientific) software. Three replicates per repeat were quantified.

### 2.2. Cell culture

#### General cell culture protocols

Human umbilical vein endothelial cells (HUVECs) were obtained from Lonza and cultured in Endothelial Growth Medium-2 (EGM-2, Lonza) or EGM-2 BulletKit (Lonza), with passages 2-6 used in experiments. Human pericytes derived from placenta were obtained from Promocell and cultured in Pericyte Growth Medium (PGM, Promocell), with passages 2-6 used in experiments. HUVECs were detached using Versene/Trypsin (9:1), while Pericytes were detached using the Detach-30 kit (Promocell).

#### Vasculogenesis cell seeding

Cells were mixed in a fibrinogen solution and injected in the device gel channel via inlets (see Figure 1A). Bovine fibrinogen (Sigma-Aldrich) and thrombin (Sigma-Aldrich) were separately dissolved in EGM-2 and DPBS, and mixed 1:1 to obtain final concentrations of 10 mg/mL and 2 U/mL, respectively. HUVECs or HUVECs/pericytes were pre-mixed in the thrombin solution to have a final cell density of 6×10^6^ HUVECs/mL, or 6×10^6^ HUVECs/mL + 6×10^5^ pericytes/mL. After injection, devices are incubated for 5 min at 37 °C. The side-channels are then filled with EGM-2 supplemented with VEGF (50 ng/mL, Peprotech) to promote vessel formation. Culture Medium is replaced every 24 h.

**Figure 1.**
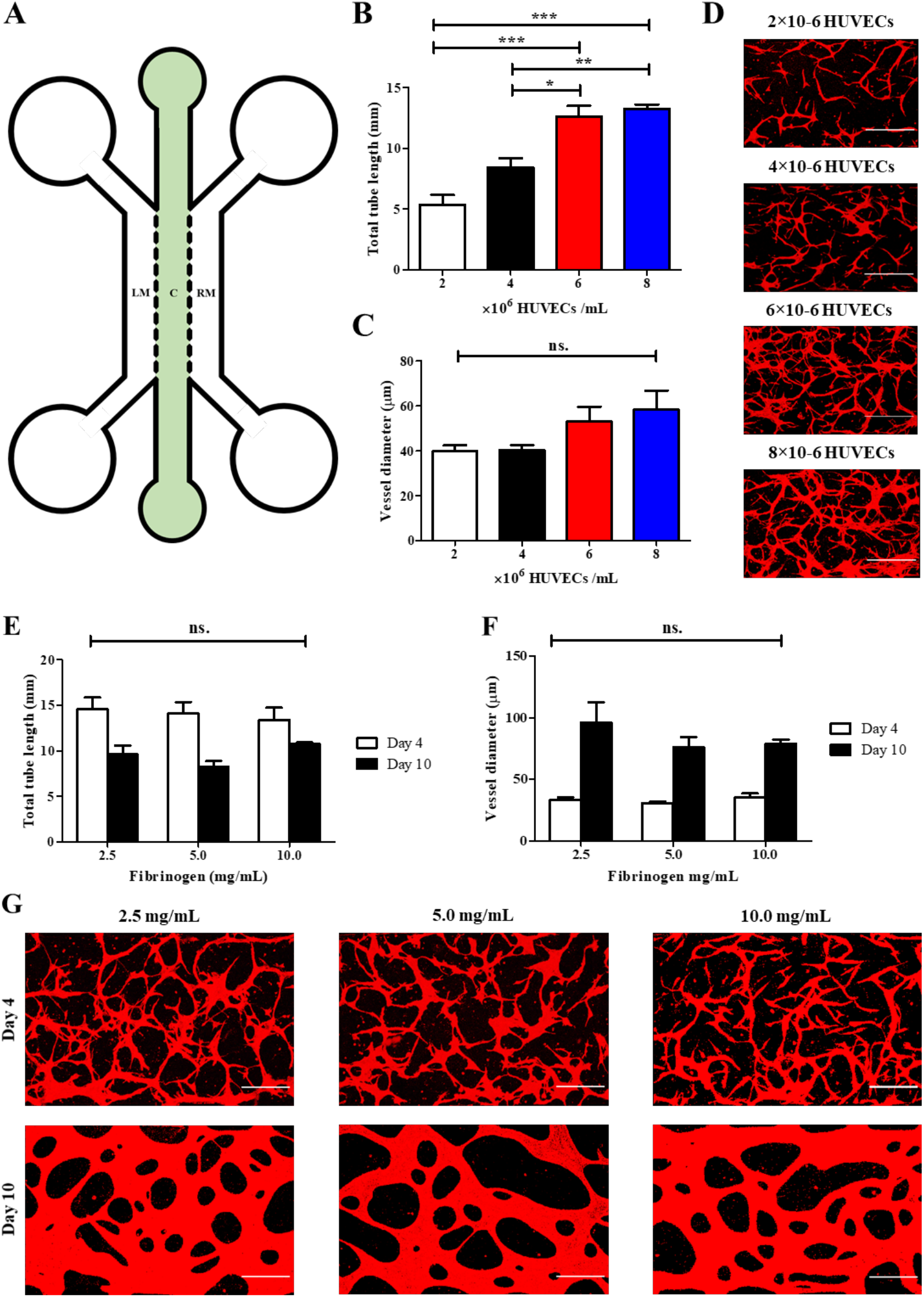
The impact of cell density and fibrinogen concentration on vasculogenesis. A) Schematic of chip design. Central channel (noted c) is 1000 µm wide and separated from the lateral medium channels (LM and RM) by 300 µm long hexagonal posts, spaced by 75 µm gaps. B) Increasing HUVEC density promotes tube formation and total tube length (5.4 ± 0.8, 8.4 ± 0.7, 12.7 ± 0.9 and 13.3 ± 0.4 mm/field of view for 2, 4, 6, 8 ×10^6^ HUVECs/mL respectively). C) HUVEC density had no significant impact on vessel diameter (mean range 39.8 - 58.4 µm). D) Representative images of different HUVEC densities, z-projection images were generated from confocal images. Red, F-actin. Scale bar: 300 µm. E) Fibrinogen concentration had no significant impact on total tube length at day 4 (mean range 13.4 - 14.5 mm/field of view), or day 10 (mean range 8.2 - 10.7 mm/field of view). F) Fibrinogen concentration also had no impact on vessel diameter following 4 days (mean range 30.6 - 35.0 µm), or 10 days of culture (mean range 75.8 - 95.8 µm). G) Representative images of networks formed at different fibrinogen concentrations, z-projection images were generated from confocal images. Red, F-actin. Scale bar: 300 µm; Statistics represents N=3.

### 2.3. Flow cytometry

We performed flow cytometry to confirm typical marker expressed by pericytes. Single cell suspensions of pericytes were stained for 30 minutes at RT using fluorescence-labelled antibodies for pericyte surface markers (488-NG2 from Invitrogen, APC PDGFR-β and APC CD105 from Biolegend) and DAPI for cell viability. Labelled cells were then washed in PBS + 1% BSA and were analyzed on a FACSCanto II flow cytometer (Becton Dickinson). Data analysis was performed using FlowJo software (Tree Star).

### 2.4. FITC-dextran permeability assay

To investigate the impact of pericytes on vascular permeability and endothelial barrier function, an assay was established based on previous reports [15, 23]. The vascular network was cultured according to the previously described protocol. Following 10-days culture, the medium reservoirs were aspirated and 30 µL EGM-2 containing 25 µg/mL 70 kDa FITC-dextran dye (Thermo Fisher Scientific) added to a single reservoir. FITC-dextran perfused through the vascular network allowing the visualisation and qualitative analysis of barrier function by recording dye diffusion across the endothelium into the extravascular compartment over a 30-minute period, using the Lumascope LS720 (Etaluma) live-imaging platform. Using ImageJ, vascular permeability was quantified using a parameter termed ‘net-fold intensity change/mm^2^’. Briefly, the intravascular and extravascular dye intensity were recorded at three regions of interest (ROI) per device (image). Following this, the change in net-fold intensity between intravascular and extravascular zones was characterised at T=0 and T=30 min - with a greater fold change indicative of more permeable vessels. T=0 was determined as when the dye intensity was stable within the vessel, therefore some devices were analysed for shorter time periods (shortest time period being 28 min). The total surface area of the vessel network was then determined using CellProfiler, by calculating the total tube length and Ferets diameter of the vessel network. Computing the ‘net-fold intensity change’ and surface area of the vessels allowed us to determine the intravascular-to-extravascular permeability of FITC-dextran, per mm^2^.

### 2.5. Nutrient starvation assay

Nutrient starvation is a well-established technique to induce cell-stress, we replicated the assay used by Nashimoto *et al* [33]. HUVEC and HUVEC/pericyte microvasculatures were cultured for 7 days. Following this, samples were cultured for a further 3 days in either normal VEGF supplemented EGM-2, or a solution of 90% DPBS and 10% VEGF supplemented EGM-2. Samples were then fixed and stained for CD31/F-actin before imaging.

### 2.6. Toxicity assay

HUVEC and HUVECs/pericytes vasculature networks were cultured for 10 days. To improve openings in the co-culture, 60 × 10^3^ HUVECs were introduced in the later media channel 24 h after seeding in the chips. The chips were then tilted at 90 degrees and incubated for 30 min. The same procedure was repeated in the other media channel. For this assay, two types of particles were used. Silica particles (Bangs Laboratories, average diameter 300 nm) were coated with poly(dimethylaminoethyl methacrylate) (PDMAEMA) following a procedure previously published [34, 35]. The final dry diameter of these particles is 330 nm. Titanium oxide (TiO_2_, anatase-SigmaAldrich) nanoparticles (diameters about 23 nm) are commercially available. Particles were washed with ethanol, centrifuged at 4k rpm for 10 min and then washed again with water and PBS with cycles of centrifugation in between. Particles were stored in PBS. Before the experiment, particles were sonicated for 5 minutes and diluted in optiMEM to prevent aggregation due to serum in the medium. The concentrations tested were: 50 and 500 µg/mL for the SiO_2_ and 100 and 1000 µM for the TiO_2_ [36]. The particles solutions were added in the media wells and cells were incubated for 4 h. Controls were also prepared with optiMEM only. After 4 h, the medium was switched back to VEGF supplemented EGM2. The samples were left for a further 4 days and then fixed and stained for CD31/F-actin.

To confirm particles entered the vasculature, tagged RNA-decorated silica particles were also prepared. Sterile particles were dispersed in PBS and mixed with an equal volume of Cy5 siRNA (Red fluorescent tagged) in RNAse free water, at an N/P ratio of 10. The mixture was then vortexed for 30 seconds and incubated at RT for 20 min. The mixture was then diluted in optiMEM and added to the media channel of the mature vasculature. After 4 h incubation, the solution was replaced with VEGF supplemented EGM2. Images using an epifluorescent microscope were acquired at several time point in fluorescent and bright field mode.

### 2.7 Immunostaining

Devices were washed with phosphate buffered saline (PBS, Sigma-Aldrich) before fixing in 4% para-formaldehyde (PFA) for 20 min at room temperature (RT). Samples were then washed with PBS and incubated with 0.4% Triton X-100 solution for 10 min at RT, before washing again with PBS. Next, samples were blocked for 4 h in 3% bovine serum albumin (BSA, Sigma-Aldrich) blocking buffer solution at RT, before overnight incubation (4 °C) with primary antibodies. The following antibodies were used: neural/glial antigen 2 (NG2) monoclonal antibody (9.2.27) Alexa Fluor 488 (eBioscience, 1:100), APC mouse anti-human CD140b (PDGFRβ) (BioLegend, 1:100), Cleaved Caspase-3 (Asp175) monoclonal antibody Alexa Fluor® 555 (Cell Signalling Technology) mouse monoclonal human CD31 Alexa Fluor 488-, 594- and 647-conjugated antibodies (BioLegend; 1:100), mouse monoclonal zona occludens-1 Alexa Fluor 594-conjugated antibody (Thermo Fisher Scientific; 1:200), mouse monoclonal beta-catenin Alexa Fluor 647-conjugated antibody (Thermo Fisher Scientific; 1:100); mouse monoclonal VE-cadherin Alexa Fluor 488-conjugated antibody (Fisher Scientific; 1:100); mouse monoclonal collagen IV (1042) Alexa Fluor 647-conjugated (eBioscience™); mouse monoclonal fibronectin (HFN7.1) Alexa Fluor 647 and 405-conjugated antibody (Novus Biological); rabbit polyclonal laminin antibody (ab11575, abcam). Cell nuclei were stained with DAPI (Sigma-Aldrich; 1:1000) and F-actin filaments were stained using phalloidin (Merck; 1:500 and Thermo Fisher Scientific; 1:40). Samples were then washed with PBS and stored at 4 °C before imaging.

### 2.8 Live/dead assay

To asses cytotoxicity after nanoparticle incubation, we used a standard LIVE/DEAD™ Viability/Cytotoxicity Kit from Thermo Fisher at the concentrations recommended by the produced. Chips were washed trice with PBS before incubation with the reagents. Samples were imaged after 30 min incubation.

### 2.9. Image analysis

F-actin or CD31 staining was used to quantify vessel formation and morphology. Following staining, vessels were imaged using the Leica TCS SP2 confocal and multiphoton microscope (Leica). Due to the 75 µm height of devices, Z-projections of the microvasculature were obtained and combined using Trans function. Following imaging, vessel formation was quantified using CellProfiler [37]. Firstly, vessel visualisation was optimised using various functions, including ‘close’ and ‘clean’, followed by skeletonization, which gave a 1-pixel wide skeleton overlay. This allowed the quantification of the overall skeleton length, which we term ‘Total Tube Length’. In addition, Feret’s diameter was determined using CellProfiler, by first quantifying the total pixelated area and dividing this by the total tube length.

Fibronectin deposition in the presence or absence of pericytes was measured by quantifying the mean intensity of the protein in the perivascular space. For this, a mask of the network was created using the CD31 staining and the intensity was recorded outside the vasculature.

For cleaved-caspase 3 characterisation and cytotoxicity analysis (via ethidium homodimer-1) after nanoparticle treatment (section 2.6), masks of the vascular network were created and the intensity of the marker was measured in the network area only. The intensity was then normalised to that measured in control samples.

### 2.10. Statistical analysis

Statistical analysis was performed on Prism (GraphPad) software. The statistical test conducted depends on the experimental paradigm and includes; unpaired one-tailed student t-test and one-way analysis of variance (ANOVA). Results are shown as mean ± standard error of the mean (SEM). Statistical significance was assumed for *p* < 0.05. * represents *p* < 0.05, ** represents *p* < 0.01, *** represents *p* < 0.001.

## 3. Results and Discussion

### 3.1. Development of vascularised microfluidic systems

Microfluidic devices (3 parallel channels separated by micro-pillars, Figure 1A) were created using photo-lithography, followed by casting of a PDMS replica and bonding to a glass coverslip. This process significantly reduces PDMS hydrophobicity and was found to induce leakage from the central compartment into the side-channels during gel loading. We investigated the recovery of hydrophobic properties of chips and observed partial recovery following storage (72 h), which was enhanced when stored at 60 °C (SI Figure S1), in agreement with the literature [38, 39]. Microfluidic devices which underwent 72 h recovery at 60^°^C displayed significant increase in the rate of successful injections, compared with chips which underwent recovery for 72 h at room temperature (100.0 ± 0.0 vs 42.9 ± 20.2%, respectively). All future devices were therefore stored for 72 h at 60 °C following plasma treatment and bonding.

Following the development of our injectable microfluidic devices, we established a basal set of conditions to form a reproducible vasculature. Initial conditions used (SI Table S1) were selected based on protocols found in the literature. We examined the impact of some of these parameters, starting with the density of endothelial cells seeded. In the literature, the endothelial cell density used to form vasculatures in a microfluidic system is in the range of 2 – 20 10^6^ HUVECs/mL [16, 24, 40]. We selected total tube length and vessel diameters to describe vessel formation following 4-days culture. As shown in Figure 1B-D, increasing HUVEC densities from 2-6 × 10^6^ HUVECs/mL led to a significant increase in total tube length, but had no significant impact on vessel diameter. In addition, no significant difference in total tube length and vessel diameter was observed between 6 and 8×10^6^ HUVECs/mL vessel networks, suggesting that 6×10^6^ HUVECs/mL is a sufficient cell density to achieve the formation of high density, relatively large vessels within our system.

Fibrinogen is the main hydrogel component in many vascularised microfluidic devices, however, the reported concentration in these studies varies substantially. The impact of this change in fibrinogen concentration is contended, with studies pointing to negative to negligible correlations with vessel formation [19, 41]. We investigated the impact of four commonly used fibrinogen concentrations (1.25, 2.5, 7.5 and 10.0 mg/mL) on vasculogenesis at days 4 and 10. HUVECs cultured in 1.25 mg/mL fibrinogen for 4 days almost entirely degraded their surrounding matrix and adhered to the underlying glass substrate (SI Figure S2A). At higher fibrinogen concentrations however, no significant impact on total tube length or vessel diameter was observed (after either 4 or 10 days of culture, see Figure 1E-G). However, there does appear to be clear morphological differences between vessels cultured for 4 or 10 days. At day 4, vessels show extensive coverage, characterised by a high total tube length (range between 11.5 - 16.1 mm/field of view) and low vessel diameter (mean range between 29-42 µm). By day 10, vessels are characterised by a low total tube length (range between 6.9 - 11.1 mm/field of view) and high vessel diameter (mean range of 65-129 µm). In addition, samples cultured for 10 days are lumenised (SI Figure S2B-C). This concurrent decrease in total tube length and increase in vessel diameter, between days 4 and 10, suggests vessels are merging to form wider overall tubes, but with a reduced overall network length. This process, reminiscent of hyperplasia, may also be associated with endothelial cell proliferation during prolonged culture times. Investigations of the impact of the presence of collagen 1 within the gel and the concentration of VEGF within the range of 25-150 ng/mL did not indicate any statistically significant difference in total tube formation (but vasculogenesis was severely restricted in the absence of VEGF, SI Figure S3-4). In addition, aprotinin treatment was only effective at limiting tube formation when supplemented in the media every 24 h. (SI Figure S5). Overall, the basal conditions we identified to form a reproducible vasculature are detailed in the Supplementary Table S1. Briefly, 6×10^6^ HUVECs/mL were cultured in a 10 mg/mL fibrinogen hydrogel for 10 days with the culture medium (EGM-2 containing 50 ng/mL VEGF) replaced every 24 h. Supplementation of fibrinogen gels with collagen or aprotinin was not continued.

### 3.2. Impact of pericytes on vessel structure

Pericytes are an important structural component of the blood vessels and the microvasculature. As such, their incorporation within forming vessel networks, in microfluidic devices, has been widely studied [15, 17, 32]. Indeed, co-culturing endothelial cells with pericytes promotes significant morphological changes in the formed vessel networks, including the inhibition of vessel hyperplasia [15].

To limit the apparent hyperplasia observed at day 10 in HUVEC monocultures, we co-cultured endothelial cells and pericytes (10:1, respectively) for 4-10 days, and quantified vessel diameter and total tube length. Similar to what is described in the literature, the introduction of pericytes led to a significant reduction in vessel diameter when compared with endothelial monocultures (36.1 ± 2.5 µm vs 52.3 ± 0.9, respectively; culture for 4 days), see Figure 2. In addition, pericytes had no significant impact on total tube length. When cultured for 10 days, co-cultures maintained a significantly lower vessel diameter, compared with endothelial monocultures (42.5 ± 2.7 vs 99.6 ± 10.8 µm, respectively). This suggests that pericytes stabilise vessel structure and prevent the occurrence of hyperplasia. In addition, a significant increase in total tube length was observed in co-cultures at day 10, compared to monocultures (13.5 ± 0.6 vs 9.5 ± 0.5 mm/field of view, respectively, Figure 2A), in agreement with the literature [17]. To confirm our conclusions, we immunostained co-cultures for the pericyte markers NG2 and PDGFR-β (Figure 2E). Cells negative for CD31 and positive for these markers can be seen spreading along the basal surface of the microvessels confirming that pericyte populations positive for PDGFR-β and NG2 associate intimately with vascular networks, recapitulating an important aspect of the normal architecture of physiological microvasculatures [29, 42].We also confirmed expression of NG2 and PDGFR-β, as well as CD105, via flow cytometry (SI Figure S6).

**Figure 2.**
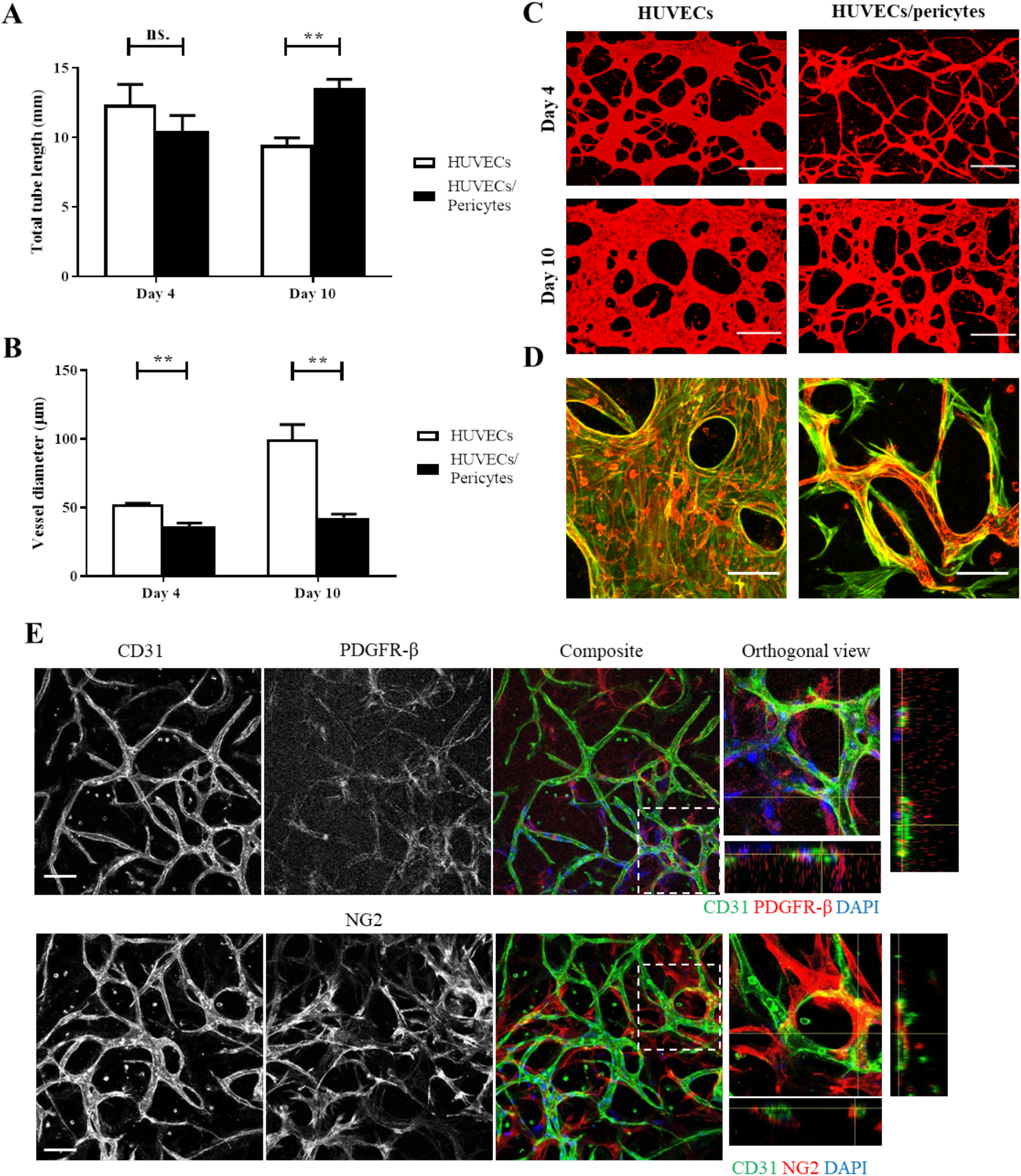
The impact of pericytes on vessel length and diameter. A) In direct co-cultures, pericytes had no significant impact on total tube length at day 4 (mean ± SEM, 12.3 ± 1.5 vs 10.4 ± 1.1, HUVECs vs HUVEC/pericytes respectively), however, a significant increase in total tube length is seen by day 10 (mean ± SEM, 9.5 ± 0.5 vs 13.5 ± 0.6, HUVECs vs HUVEC/pericytes respectively). B) The addition of pericytes also led to a significant reduction in vessel diameter following both 4 and 10 days of culture. C) Representative images of samples, z-projection images were generated using confocal images. Red, CD31. Scale bar: 300 µm. D) Representative images showing HUVEC-pericyte interactions, z-projection images were generated using confocal images. Red, CD31. Green, F-actin. Scale bar: 100 µm. Statistics represents N=3. E) Cells expressing the pericyte-specific markers PDGFR-β and NG2 (red) are clearly seen at the surface of the microvascular network. Corresponding orthogonal views indicate that these cells are often found wrapping around CD31 positive vessels. Scale bar: 100 μm.

To investigate paracrine signalling between pericytes and endothelial cells we developed an alternative chip design incorporating two parallel gel channels (SI Figure S7). Seeding HUVECs and pericytes in separate parallel compartments, we examined if pericyte paracrine signalling impacted vascular network morphology (no exogenous VEGF was added). Following 4-days culture, the addition of pericytes led to an increase in total tube length (mean ± SEM, 3.2 ± 0.2 vs 1.4 ± 0.3 mm/field of view), however had no significant impact on vessel diameter. Hence, although some level of paracrine signalling may result in increased stability of vessels and promote tube formation, results were not as marked as in direct co-culture experiments.

### 3.3. Impact of pericytes on vessel maturation

In addition to regulating morphological features of the endothelium, pericytes also promote vessel maturity and barrier function [43-45]. To indicate vessel maturation, the expression of several cell-cell junction proteins, notably members of the tight and adherens junctions, are often characterised [15, 22, 23, 46]. We examined the recruitment of tight junction protein ZO-1, and cell adhesion proteins VE-cadherin and β-catenin at cell-cell junctions (see Figure 3A and SI Figure S8). Confocal images did not indicate any significant changes in recruitment of these junction markers in co-cultures compared to monocultures. In particular, ZO-1 was clearly recruited at cell-cell junctions, with little diffuse background staining, suggesting that endothelial cells in both conditions form mature endothelial barriers. Similar recruitment patterns were observed for VE-cadherin and β-catenin. Expression of these tight and adherens junction proteins are routinely demonstrated to indicate vessels which have reached maturation [15, 16, 22, 23, 46]. Accordingly, in our system endothelial cells have differentiated into mature vessels following 10-days culture. In addition, endothelial cells typically displayed an elongated phenotype aligned along the length of the vessels. However, in wider vessels some endothelial cells displayed a more cuboidal morphology, with reduced alignment, in particular in endothelial monocultures.

**Figure 3.**
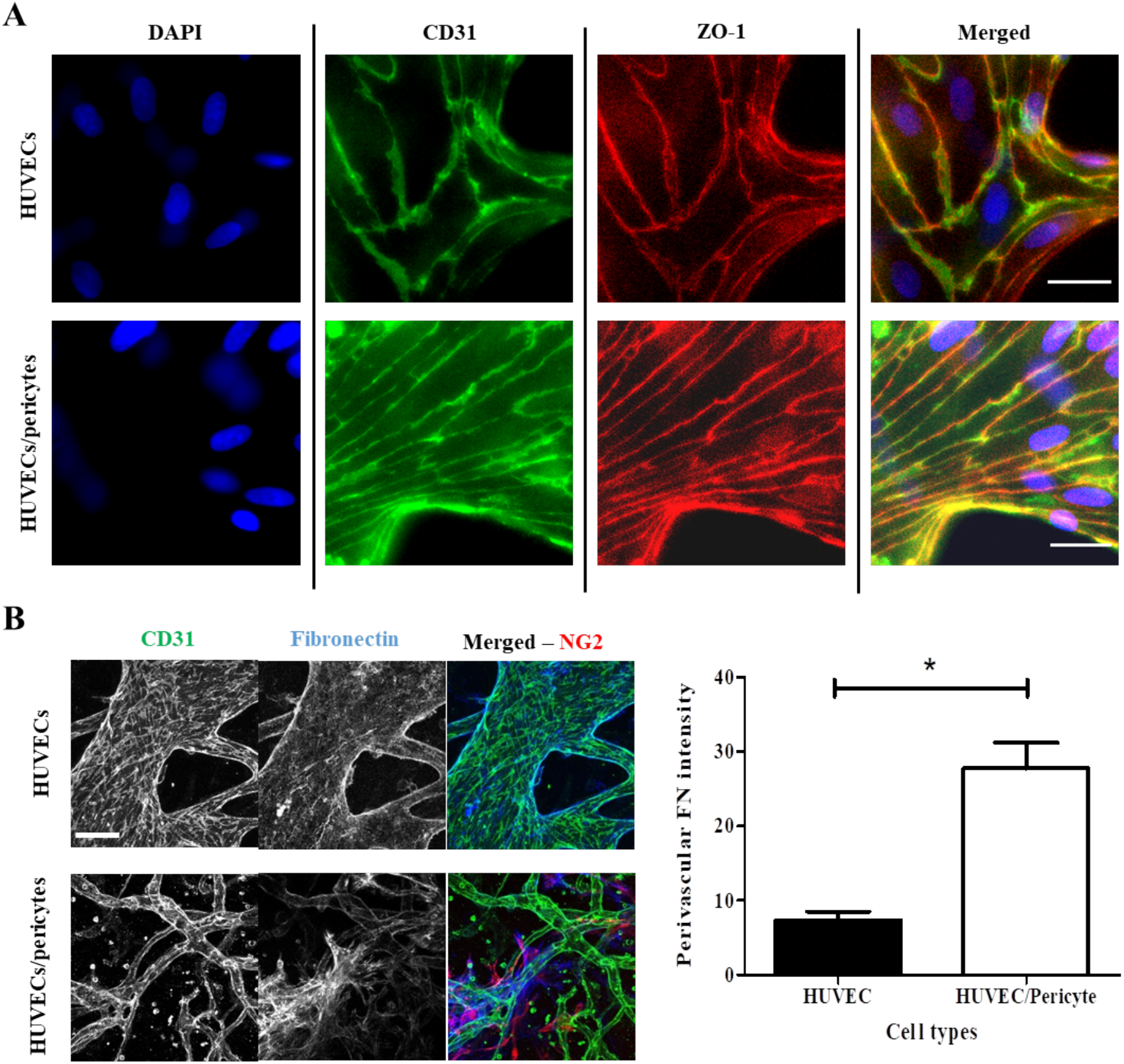
Vessel maturation markers. A) Epifluorescence microscopy images of HUVECs or HUVECs/pericytes vessel networks following 10-days culture, z-projection images were generated using epifluorescence images. These images represent a single Z-frame, displaying junction expression of CD31 and ZO-1. Blue, DAPI. Green, CD31. Red, ZO-1. Scale bar: 25 µm. B) Images displaying fibronectin deposition (z-projection images were generated via confocal microscopy). Blue, Fibronectin. Green, CD31. Red, NG2. Scale bar: 100 µm. C) Quantification of fibronectin deposited in the perivascular space.

The basement membrane has an important physiological role in the morphology and development of blood vessels and abnormalities have been reported in tumour angiogenesis [47, 48]. The presence of pericytes has been shown to regulate the deposition of ECM proteins at the basement membrane [49]. Type IV collagen and laminin are two of the main components of the basement membrane, whereas fibronectin is deposited by fibroblasts and pericytes in the surrounding mesenchymal tissue. We assessed the deposition of these three proteins in mature vasculatures (after 10-days culture) in mono- and co-cultures. Images show that all three proteins were secreted in both conditions and are tightly associated with the vascular network (Figure 3B and SI Figure S9). Orthogonal optical sections obtained for collagen IV and laminin indicated proteins deposition at the basement membrane of endothelial networks, in both mono- and co-cultures. However, fibronectin deposition accumulated in interstitial tissue in co-cultures, closely associated with pericytes. Quantification of the perivascular fibronectin, deposited adjacent to the vascular network, but not associated with its lamina, confirmed significant levels of fibronectin deposition in the perivascular space in the presence of pericytes, compared to monocultures (Figure 3B).

Following culture for 10 days, we demonstrated vessels display both junction marker expression and localisation, and ECM deposition associated with mature microvasculature formation (Figure 3). Therefore, we next investigated the impact of pericytes on vessel permeability in our system at this time point. Using protocols comparable to those reported by others, we perfused 70 kDa FITC-dextran through the vessel network [15, 21, 23, 33, 50]. Quantitative analysis of diffusion to interstitial gel areas indicated no change significant difference in the cross-endothelial diffusion of this dye, when comparing HUVEC monocultures and HUVEC/ pericyte co-cultures (Figure 4). Stronger differences in diffusion barrier were reported in the literature, indicative of pericytes contributing to the maturation of the endothelial barrier [17, 51]. However, the moderate impact that pericytes had on barrier function is in line with the quality of cell-cell junctions and the basement membrane observed in both mono- and co-cultures.

**Figure 4.**
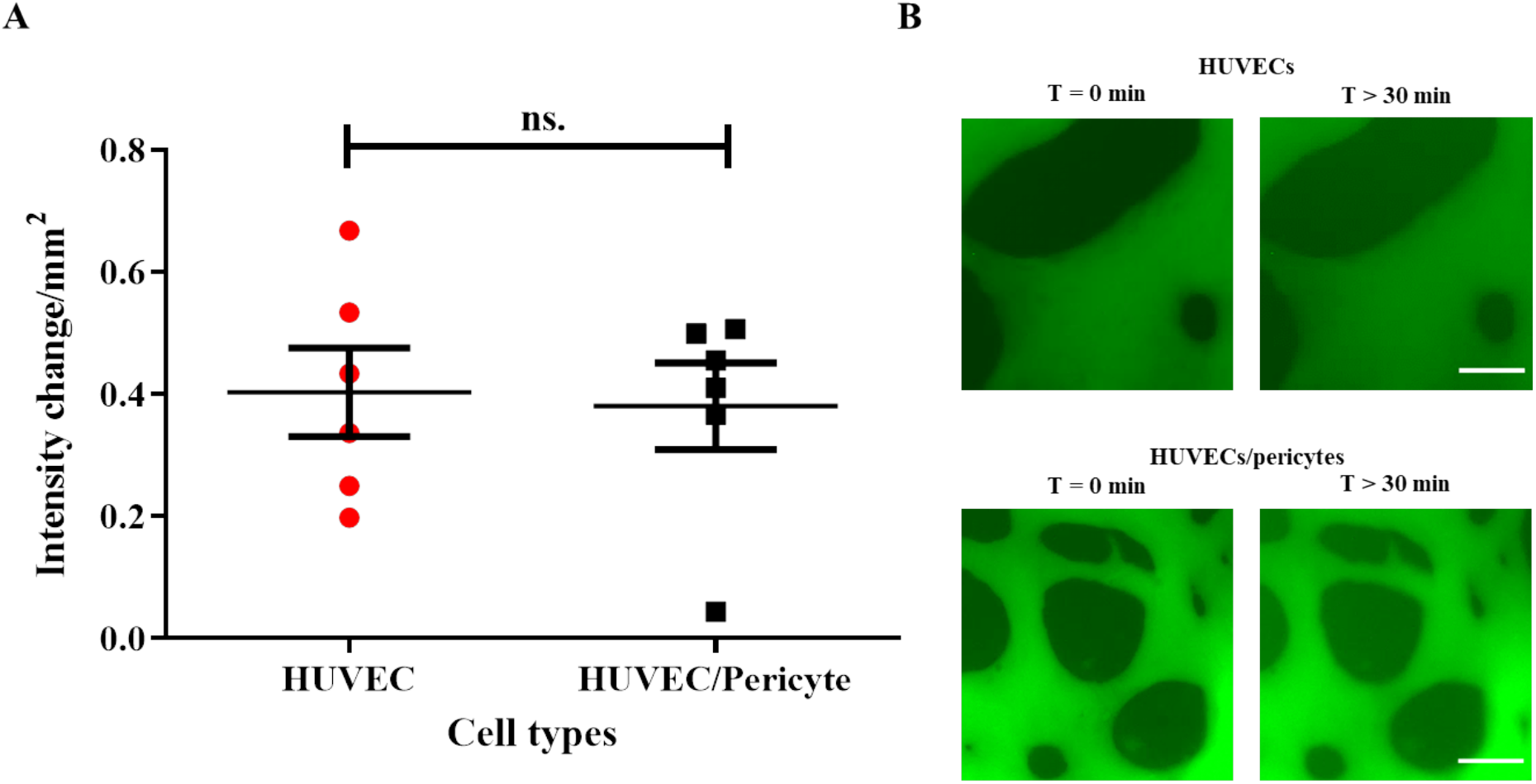
The impact of pericytes on vessel permeability. 70 kDa FITC-dextran was perfused after 10 days of culture. A.) Using ‘net-fold intensity change/mm^2^ to define vessel permeability, the addition of pericytes had no significant impact on endothelial barrier function (mean ± SEM, 0.38 ± 0.07 compared with 0.4 ± 0.07 for endothelial monocultures). B.) Representative epifluorescence images at T = 0 and T = 30 min. Scale bar: 100 µm. Statistics represents N=6, 3 ROI per 2 independent repeats.

Therefore, overall, our results indicate that pericytes play a minor role in the development and maintenance of the barrier function of microvascularised networks generated in microfluidic chips. Although significant changes in network morphology are observed in co-cultures, cell-cell junctions appear well established and their impact on barrier function is broadly maintained. However, pericytes had a striking impact on the perivascular matrix remodelling, rather than basement membrane remodelling per se, therefore suggesting that they may play an important role in the stabilisation of vascular networks.

### 3.4. Impact of pericytes on microvasculature stability

Pericytes are important regulators of endothelial survival and vessel integrity [30, 31, 52]. To investigate this further we compared the impact of nutrient starvation on endothelial monocultures and pericyte co-cultures (Figure 5) using a method adopted from Nashimoto *et al* [33]. Serum and nutrient starvation assays are commonly used to investigate environmental stress and apoptosis and have been shown to induce endothelial cell death [53-55]. Following 72 h nutrient deprivation in endothelial monoculture vessel networks, cell-cell adhesion was severely compromised and HUVECs formed small cell aggregates (Figure 5C), indicative of apoptosis [56]. Interestingly, under identical conditions, co-culture networks retained their structure, although a reduction in total tube length was observed, compared to untreated controls (13.5 ± 0.6 vs 10.7 ± 0.7 mm/field of view, respectively). In addition, nutrient depravation had no significant impact on co-culture vessel diameter (Figure 5B). Some single rounded cells were observed in nutrient starved co-cultures, similar to those observed in HUVEC monocultures, although these were much less prevalent. Therefore, our data suggests that pericytes promote the stability and integrity of endothelial networks, even under severe stress conditions. This agrees with observations that pericytes play a role in the survival of endothelial cells [30, 31, 52].

**Figure 5.**
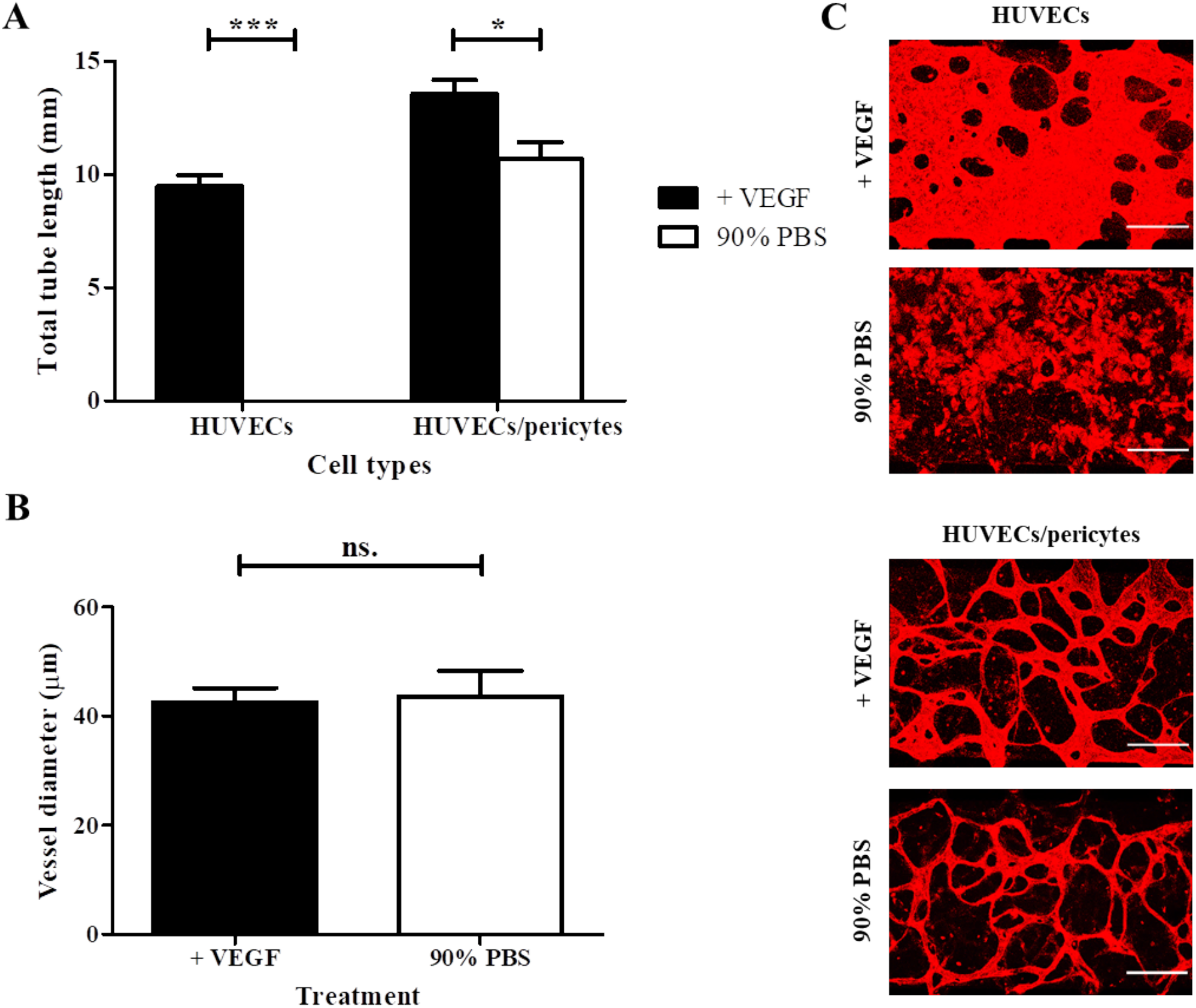
The impact of pericytes on network stability in response to starvation. A) 72 h nutrient starvation resulted in the entire collapse of HUVEC networks in monocultures. In contrast, HUVEC/ pericyte networks persisted, although they apparently regressed compared with untreated conditions (mean ± SEM total tube length, 10.7 ± 0.7 vs 13.5 ± 0.6 mm/field of view, respectively). B) Impact of starvation on tube diameter in co-cultures, showing no significant difference after starvation.C) Representative images, z-projection images were generated using confocal images. Red, CD31. Scale bar: 300 µm. Statistics represents N=3.

Considering the impact of pericytes on the stability of networks in response to stress, we proposed that they could play a role in the preservation of network stability in response to toxic compounds such as some nanoparticles. TiO_2_ nanoparticles displayed mild toxicity on 2D endothelial layers, disrupting cell-cell junctions and inducing leakiness [36]. We investigated the impact of these nanomaterials (TiO_2_ nanoparticles with a diameter of 23 nm), as well as that of other positively charged polymer-coated silica nanoparticles (330 nm diameter with a poly(dimethyl aminoethyl methacrylate) shell). These latter particles are finding applications in gene delivery and known to induce toxicity in endothelial cells [35]. 100 and 1000 μM TiO_2_ nanoparticles displayed a negligible impact on total tube length and vessel diameter in both the mono and co-cultures (SI Figure S10), in contrast to their impact on endothelial monolayers (tube diameters of 137.3 ± 33.1, 106.5 ± 24.4 and 90.2 ± 7.3 μm for the control and 100 and 1000 μM TiO_2_ exposed samples, respectively), although this is not significant [36]. Cationic PDMAEMA-coated silica nanoparticles displayed a significant effect on the HUVECs microvasculature at both 50 and 500 µg/mL concentrations, destabilising associated networks at the highest concentrations tested (Figure 6). While the total tube length was not significantly affected, the tube diameter decreased from 137.3 ± 33.1 μm for control to 76.2 ± 13.5 and 49.6 ± 10.6 μm for the 50 and 500 μg/mL, respectively. Associated with these morphological changes, the density of network branches was not significantly affected in co-cultures but clearly increased in the HUVECs networks, reflecting the gradual breakdown of corresponding networks (SI Figure S11). Overall, images clearly evidenced the disassembly of the networks following exposure to these nanoparticles. However, co-cultured networks remained stable even at the highest concentrations of nanoparticles tested, with no significant impact on total tube length and vessel diameter (Figure 6, tube diameter is 29.2 ± 2.0 and 24.8 ± 2.3 μm for control and 500 μg/mL particles respectively).

**Figure 6.**
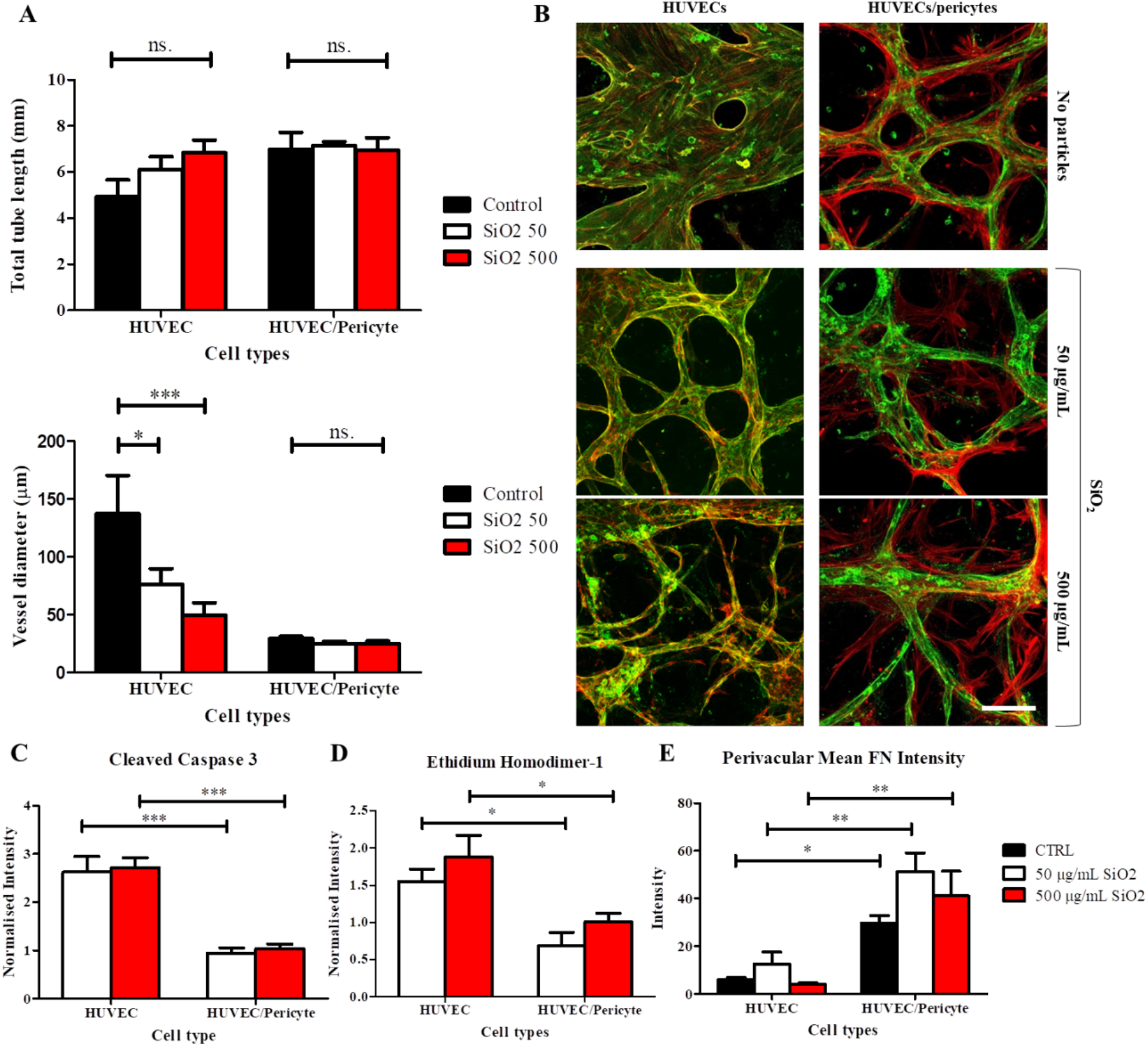
The impact of cationic nanoparticles on vascular network integrity. A) Following 4-day treatment with cationic silica nanoparticles, a significant reduction in vessel diameter is observed in HUVEC monocultures. This is not observed when HUVECs are co-cultured with pericytes. B) Representative images of microvascular networks following treatment with cationic silica nanoparticles, z-projection images were generated using confocal images. Red, phalloidin. Green, CD31. Scale bar: 100 μm. C-E) Quantification of relative mean intensity for cleaved caspase 3 (ratio treated/Ctrl) and cytotoxicity via ethidium homodimer-1 (ratio treated/Ctrl) and mean intensity of perivascular fibronectin (FN) deposition. Statistics represent N=3.

To investigate processes associated with this response to cationic nanoparticles, we assessed apoptosis, through cleaved caspase 3 expression and ethidium homodimer-1 nuclear localisation (Figure 6C-E and SI Figure S12). Cleaved caspase 3 expression was significantly higher in nanoparticle-treated monocultures compared to untreated controls (realtive intensity ratios of 2.6 ± 0.3 and 2.7 ± 0.2 at nanoparticle concentrations of 50 and 500 μg/mL, respectively; Figure 6C). In contrast, cleaved caspase 3 expression in co-culture remained comparable to controls (ratios are 0.9 ± 0.1 and 1 ± 0.09 at nanoparticle concentrations of 50 and 500 μg/mL, respectively). Similarly ethidium homodimer-1 nuclear localisation increased in monocultures treated with cationic nanoparticles treatment, but not in co-cultures (Figure 6D). Therefore, our data indicate a protective role of pericytes on nanoparticle-induced apoptosis.

Perivascular fibronectin deposition in response to cationic nanoparticle treatment was next examined (Figure 6E). Fibronectin deposition was unaltered by nanoparticle exposure in both mono- and co-cultures treated with cationic nanoparticle, compared to untreated controls, but was found to be systematically higher in co-cultures, even after cationic nanoparticles treatment. Hence the increased fibronectin deposition observed in co-cultures was preserved in the context of a nanotoxicity response.

Finally, we confirmed that nanoparticles accumulated in these networks after several hours of treatment, using silica particles loaded with fluorescently labelled RNA (SI Figure S10). Therefore, our results indicate that, in contrast to the weak impact of the negatively charged TiO_2_ nanoparticles, cationic silica nanoparticles resulted in significant cell toxicity, even in a 3D model. However, despite the significant stress induced to the network, pericyte co-culture stabilised associated microvasculatures.

In addition to their anti-apoptotic paracrine effect [30, 31], pericytes may contribute to the stability of microvascular networks via other mechanisms. Our results clearly indicate an increased remodelling of the perivascular matrix, with significant fibronectin deposition, which may contribute to the stability of the apico-basal polarity in response to environmental stress, such as in serum starvation and in response to cationic nanoparticles. It could also be proposed that other structures associated with vasculature maturity, such as the glycocalyx composition and surface density at the membrane [57], contribute to confer protection from contact toxicity (for example associated with cationic nanoparticles), without significantly regulating barrier function. Further studies are required in order to conclude on the precise mechanism via which pericytes stabilise vascular networks in response to stress and nanostoxicity in vitro and in vivo.

## 4. Conclusion

With our model, we quantified the impact of a range of parameters involved in the establishment of a microvasculature in multi-channel microfluidic chips. Our work highlighted the role of pericytes in the stabilisation of vascular networks and the prevention of hyperplasia. We demonstrated that pericytes, through paracrine signalling, promote short-term vessel formation. However, our results indicate that it is perivascular matrix remodelling that regulates vessel hyperplasia and contribute to the stabilisation of the vessel plexus. Although we found pericytes had no significant impact on barrier function, we found that pericytes protected the integrity of vascular networks and inhibited vessel regression following nutrient starvation when compared with endothelial monocultures. Furthermore, a similar protective response was observed in co-cultures exposed to high concentrations of cationic nanoparticles. These results highlight the importance of pericytes in protecting the microvasculature, and clearly demonstrate that *in vitro* models which feature microvasculature stress should incorporate pericytes, to capture their protective effect on endothelial integrity, whether for safety and toxicity testing or to model the progression of diseases.

## Acknowldegments

Funding from the National Centre for the Replacement, Refinement and Reduction of Animals in Research (NC3Rs, NC/M001636/1), Cancer Research UK (Programme Grants C587/A16354 and C587/A25714) and from the European Research Council (ProLiCell, 772462 and CANBUILD, 32566) is gratefully acknowledged.

