## Supplementary Information for "Impact of Pericytes on the Stabilisation of Microvascular Networks in Microfluidic Systems in Response to Nanotoxicity"

**Table S1. Standard fibrinogen gel components**

| **Component** | ***Initial*** **Concentration/time** | ***Finalised*** **Concentration/time** |
|---|---|---|
| **HUVECs (10-6/mL)** | 6 | 6 |
| **Fibrinogen (mg/mL)** | 2.5 (PBS) | 10 (PBS) |
| **Thrombin (U/mL)** | 2 (PBS) | 2 (EGM-2) |
| **Type 1 Collagen (mg/mL)** | 0.2 | - |
| **Aprotinin (U/mL)** | 0.15 | - |
| **VEGF (ng/mL)** | 50 | 50 |
| **Duration (Days)** | 4 | 10 |

Initial conditions were selected according to typical procedures from the literature. After evaluation of different parameters, we developed a finalised basal set of conditions. Aprotinin and Collagen 1 were not found to significantly impact the quality of networks generated.

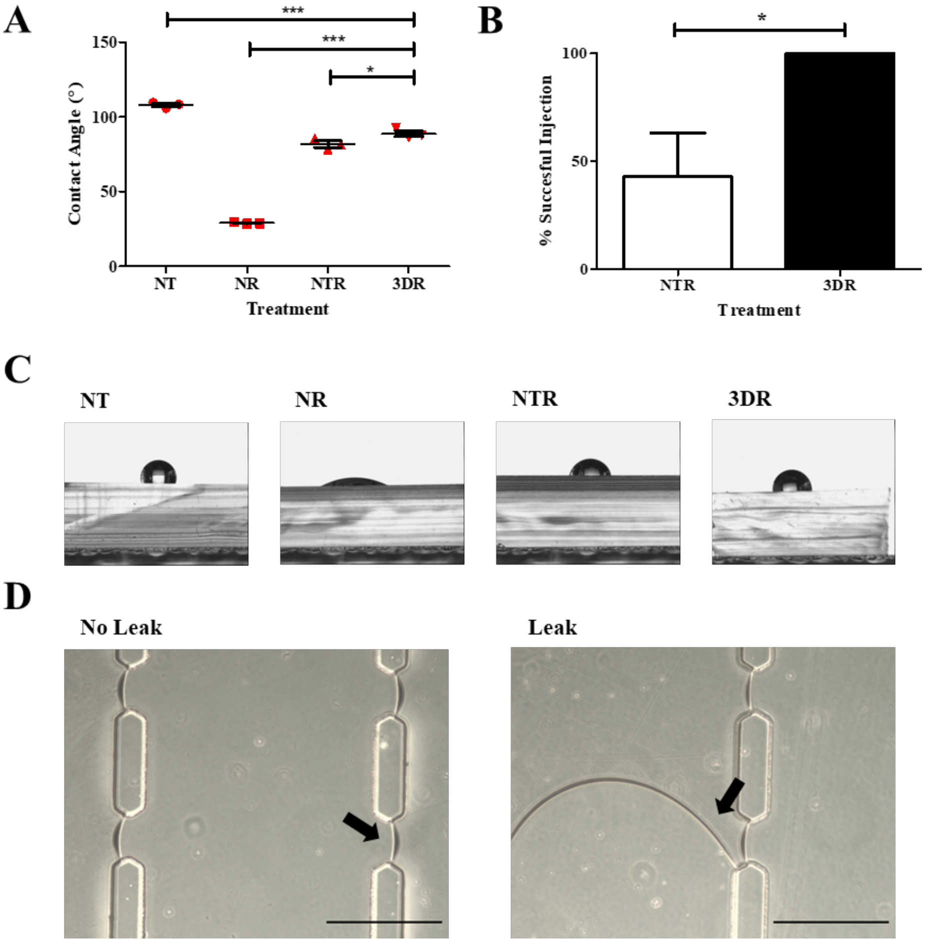


**Figure S1. Processing microfluidic devices.** A) Characterisation of hydrophobicity of PDMS following processing steps typical of microfluidic chip fabrication. Acronyms: NT - No Treatment, NR - No Recovery, NTR - No Thermal Recovery, HR - Heated Recovery (60 °C). Statistics correspond to N=3. B) Restoration of hydrophobicity through thermal recovery also led to a significant increase in successful injections when compared with NTR samples. Statistics correspond to 15 samples. C) Representative images of water droplets at PDMS interfaces tested in A. D) Representative images of gel injection, with and without gel leakage. The corresponding interfaces are indicated by arrows. Scale bar: 300 µm.

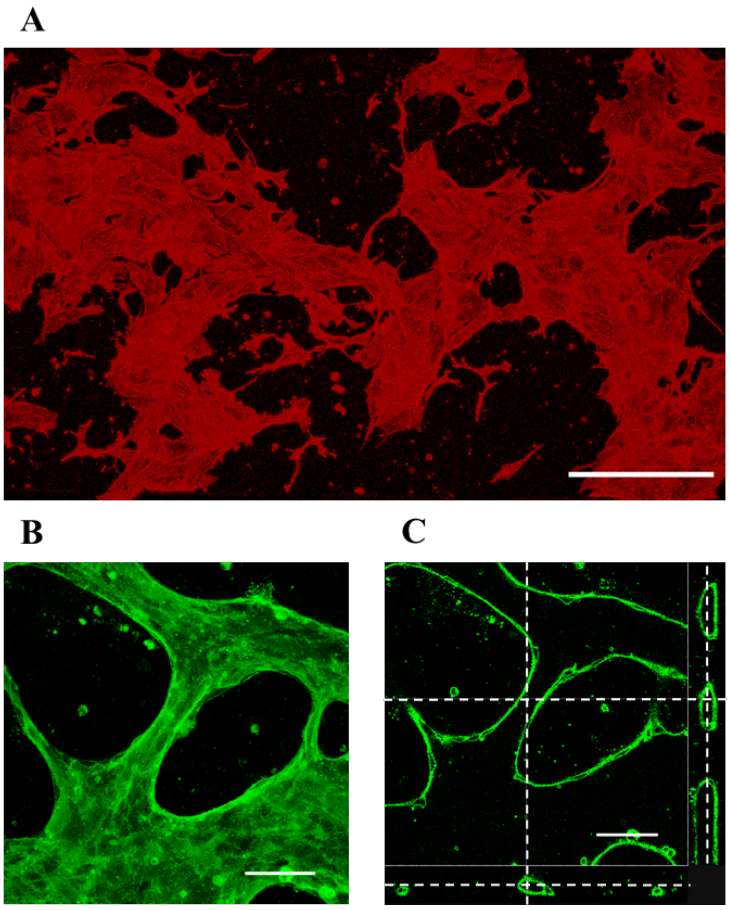


**Figure S2. Representative images HUVEC networks in monoculture.** A) Following 4-days culture in 1.25 mg/mL fibrinogen, HUVECs had extensively degraded the surrounding ECM and adhered to the underlying glass substrate. Red, F-actin. Scale bar: 300 µm. B) Z-stack of microvascularised networks following 10-day monocultures in 10 mg/mL fibrinogen. Green, CD31. Scale bar: 75 µm. C) Cross-section of B clearly demonstrating the formation of lumenised vessels. Green, CD31. Scale bar: 75 µm.

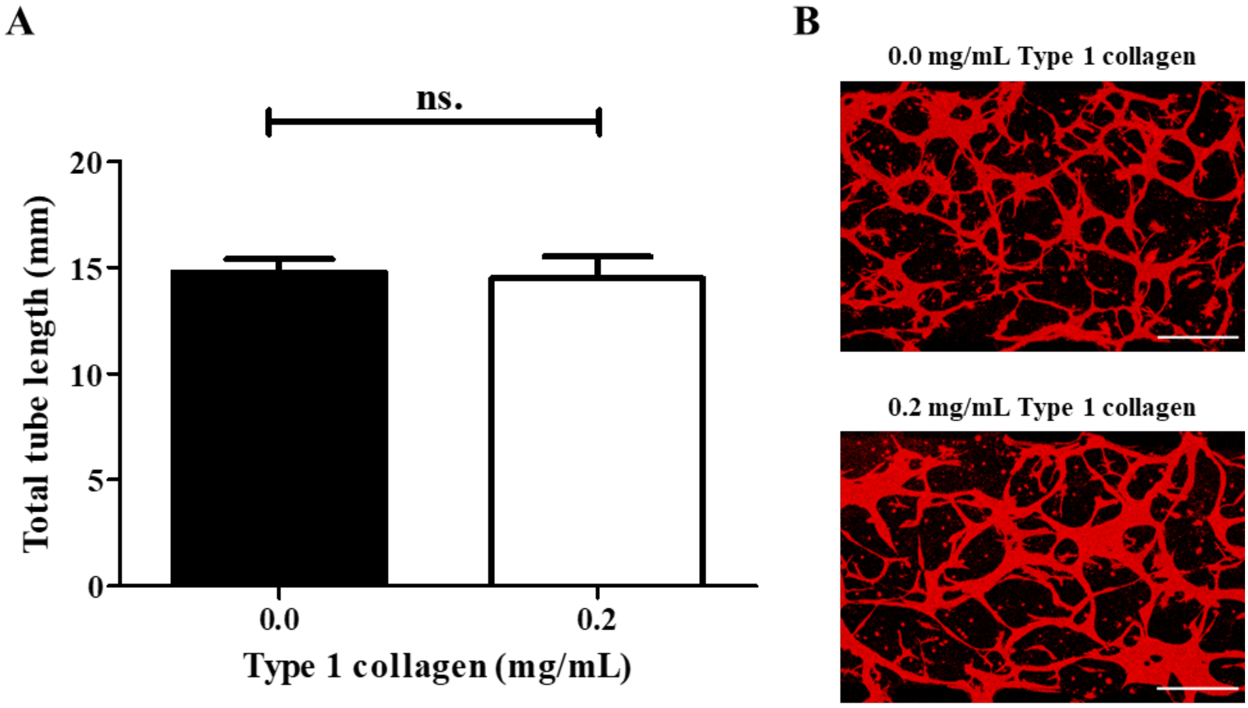


**Figure S3. Impact of type 1 collagen on vessel formation.** A) Addition of 0.2 mg/mL type 1 collagen to fibrinogen gels has no significant impact on total tube length compared with control, 14.5 ± 1.0 vs 14.8 ± 0.6 mm/field of view, respectively. Samples were cultured for 4 days, in the presence of 50 ng/mL VEGF. B) Representative images. Red, phalloidin. Scale bar: 300 µm. Statistics correspond to N=3.

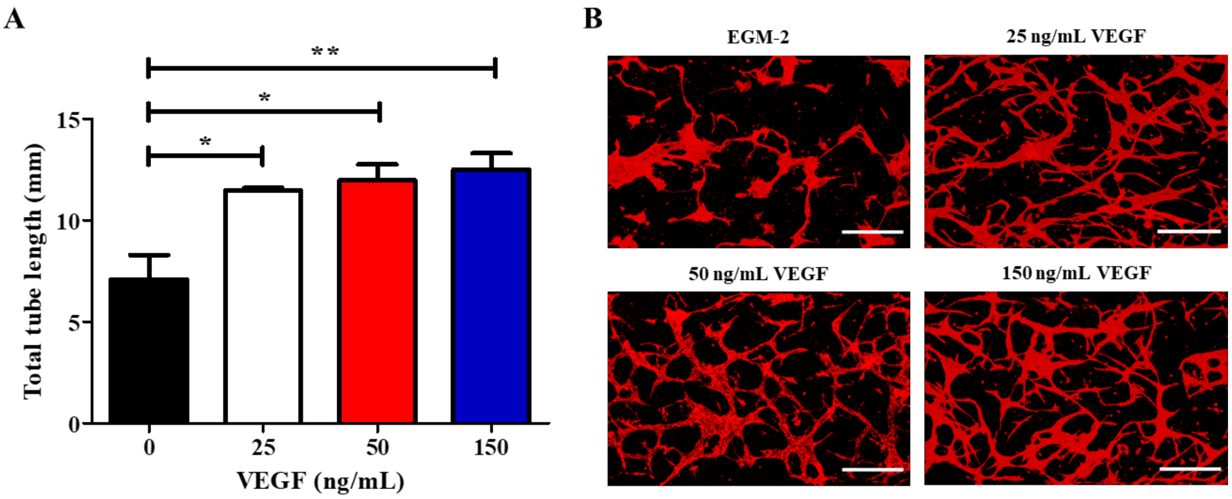


**Figure S4. Impact of VEGF on vessel formation.** A) Impact of VEGF was tested at 3 different concentrations (25, 50 and 150 ng/mL) and a negative control. All VEGF concentrations promoted a significant increase in total tube length compared with the control (mean ± SEM, 7.1 ± 1.2, 11.5 ± 0.1, 12.0 ± 0.8 and 12.5 ± 0.8 mm/field of view, for 0, 25, 50 and 150 ng/m/ VEGF, respectively). No significant difference in tube formation is observed between any of the VEGF containing groups tested. Samples were cultured for 4 days. B) Representative images. Red, phalloidin. Scale bar: 300 µm. Statistics correspond to N=3.

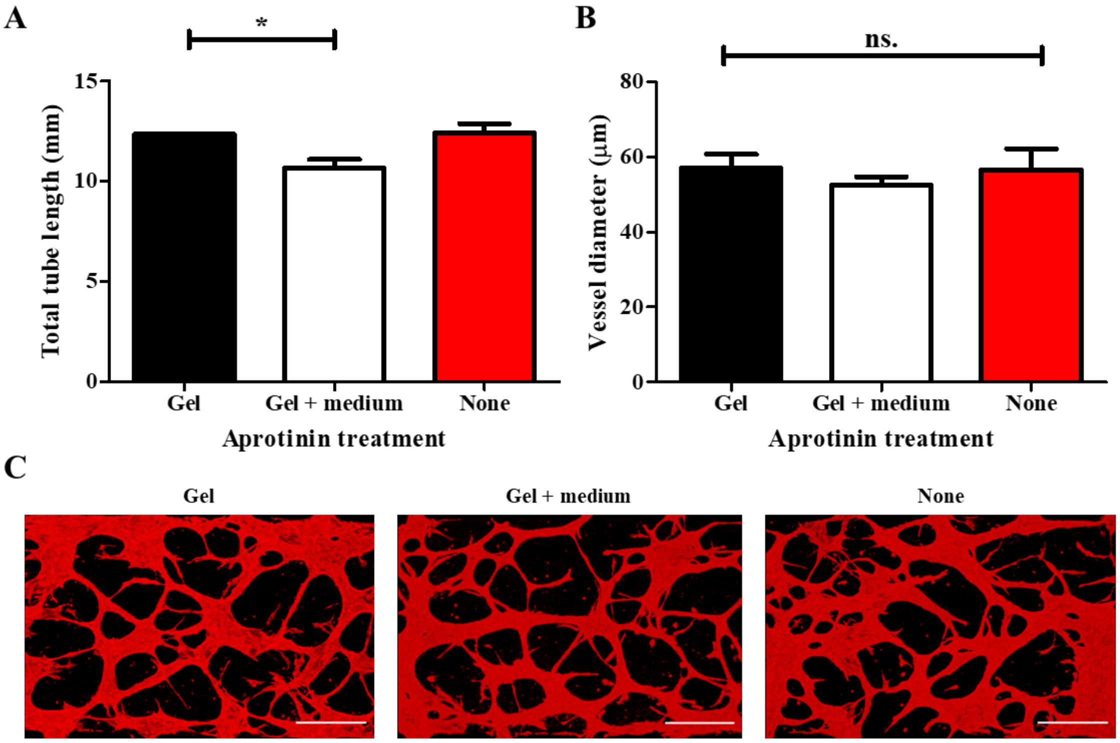


**Figure S5. Impact of aprotinin on vessel formation.** Three different conditions were investigated: gel, 0.15 U/mL aprotinin supplemented during fibrin gel formation alone; gel + medium, 0.15 U/mL aprotinin supplemented throughout; None, no aprotinin supplemented at any point. Samples were cultured for 4 days, in the presence of 50 ng/mL VEGF. A) Supplementing the gel + medium with aprotinin led to a loss in total tube length when compared with only supplementing the gel (mean ± SEM, 12.3 ± 0.0 vs 10.65 ± 0.4 mm/field of view, respectively). However, there is no significant difference in total tube length between gel and no aprotinin. B) Aprotinin has no significant impact on vessel diameter. Vessel diameters: 57.0 ± 3.8, 52.4 ± 2.3 and 56.6 ± 5.5 µm for control, + and - aprotinin, respectively. C) Representative images. Red, phalloidin. Scale bar: 300 µm. Statistics correspond to N=4.

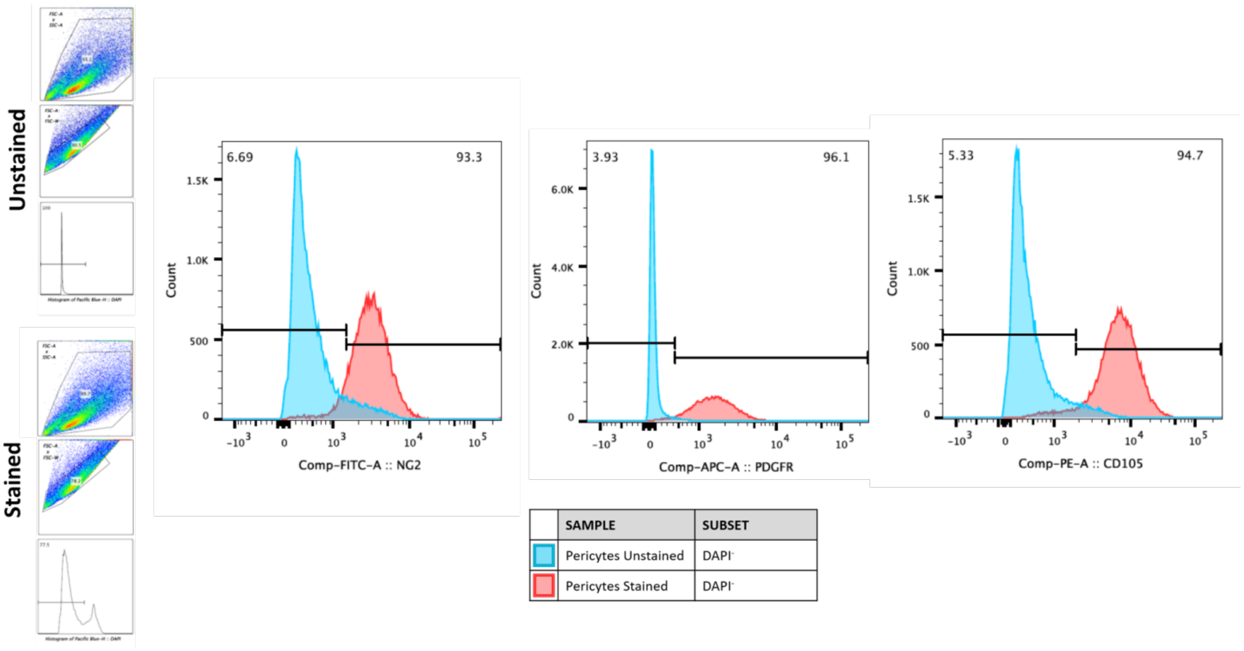


**Figure S6. Pericytes flow cytometry.** Expression of NG2, PDGFR-β and CD105 in pericytes was assessed via flow cytometry.

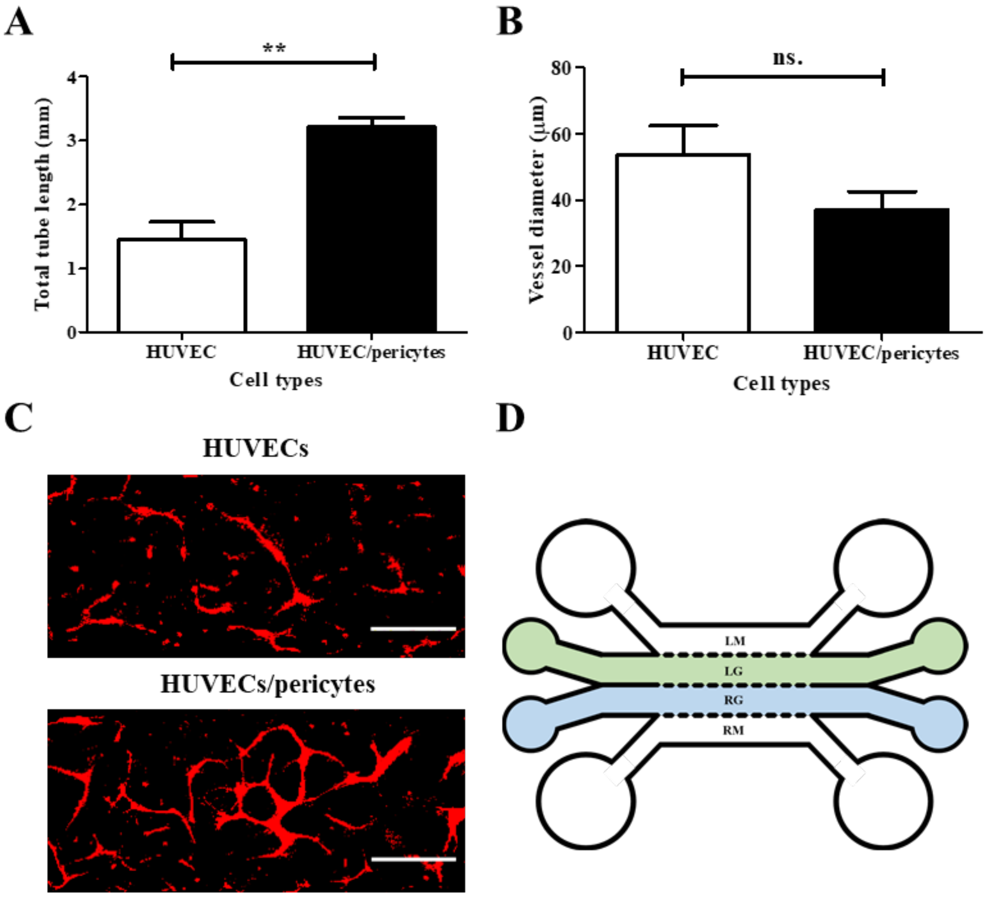


**Figure S7. Investigation of pericyte-HUVEC paracrine signalling.** The impact of pericyte paracrine signalling on total tube formation in the absence of exogenous VEGF, following 4-day culture was examined. Pericytes and HUVECs were seeded in separate, parallel gel channels. A) The addition of pericytes promoted a significant increase in total tube formation compared with negative control (mean ± SEM, 3.2 ± 0.2 vs 1.4 ± 0.3 mm/field of view, respectively). B) However, though there is a drop in vessel diameter with the addition of pericytes (mean ± SEM 37.0 ± 5.6 mm/field of view) compared with HUVEC mono-culture ((mean ± SEM 53.6 ± 8.8 mm/field of view), it is not statistically significant. C) Representative images. Red, CD31. Scale bar 300: µm. D) Schematic of corresponding chip design. Central gel channels (LG and RG) are 700 µm wide and separated from parallel channels by 100 µm long trapezoidal posts, spaced 100 µm.

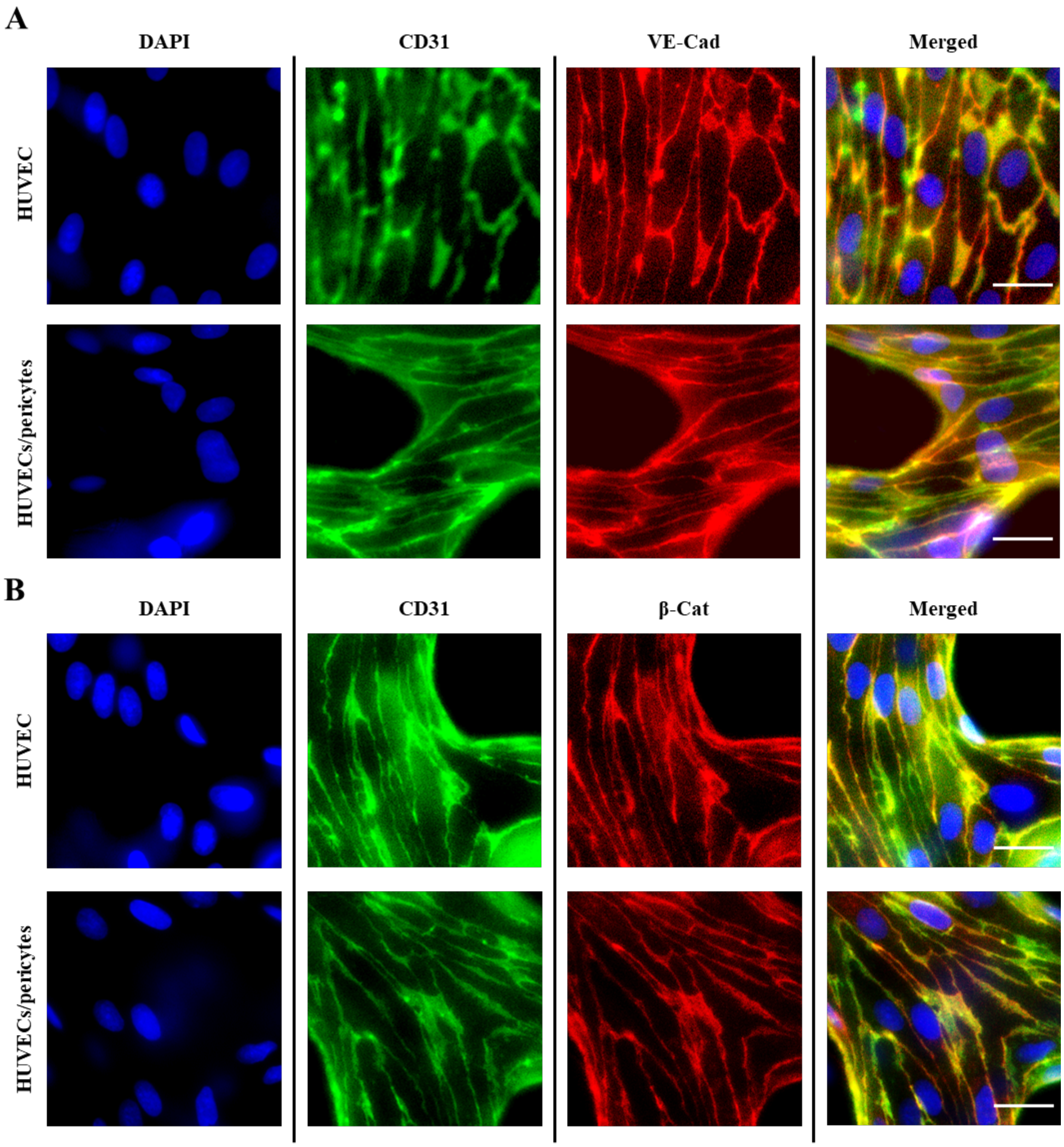


**Figure S8. Characterisation of endothelial junction markers.** A) Epifluorescence microscopy images of HUVECs or HUVECs/ pericytes vessel networks following 10-day culture. These images represent a single Z-frame, displaying junction expression of CD31 and VE-cadherin. Blue, DAPI. Green, CD31. Red, VE-Cad. B) These vessels display junction expression of CD31 and β-catenin. Blue, DAPI. Green, CD31. Red, β-cat. Scale bar: 25 μm.

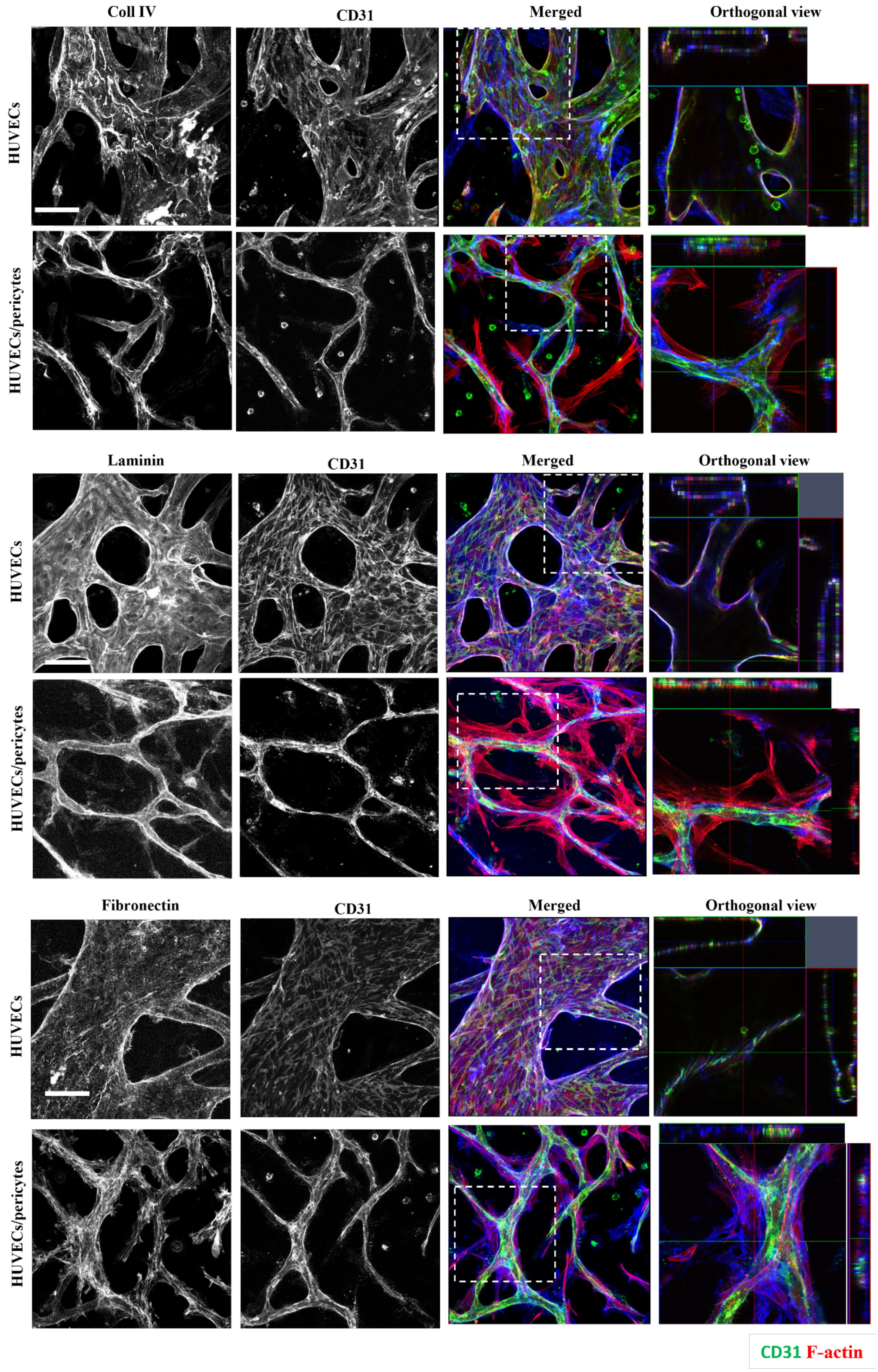
Coll IV
CD31
Merged
Orthogonal view
HUVECs
HUVECs/pericytes
Laminin
CD31
Merged
Orthogonal view
HUVECs
HUVECs/pericytes
Fibronectin
CD31
Merged
Orthogonal view
HUVECs
HUVECs/pericytes
CD31 F-actin

**Figure S9. Matrix deposition during network assembly.** Confocal images (z-projections) of ECM protein deposition: collagen IV, fibronectin and laminin. Proteins are tightly associated with the vasculature, but fibronectin images also indicate some deposition in the perivascular space, as can be seen in the orthogonal view. Blue, ECM protein. Green, CD31. Red, F-actin. Scale bar: 100 μm.

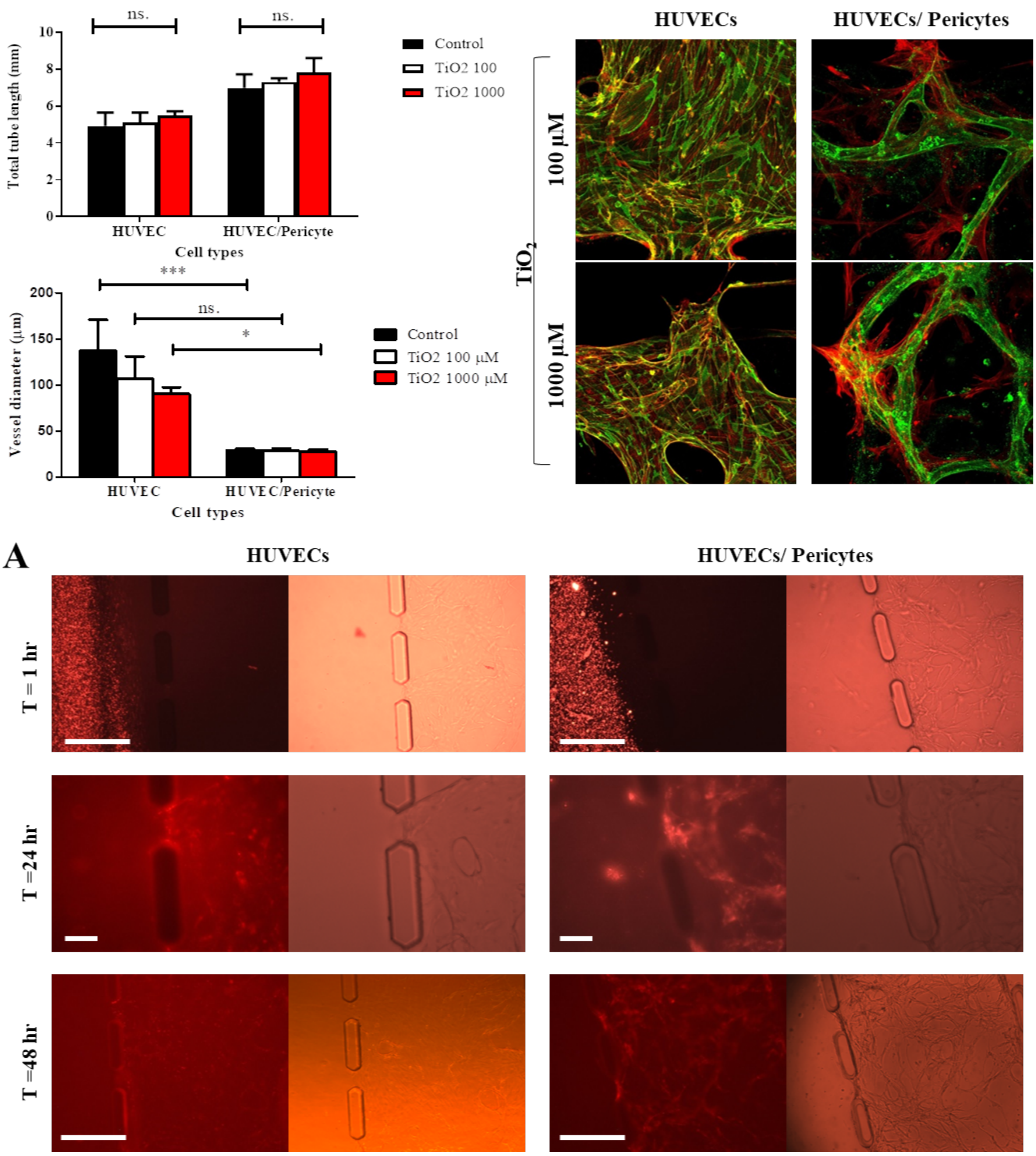


**Figure S10. Impact of $TiO_2$ nanoparticles on vascular integrity.** A) Confocal images of HUVECs and HUVECs/Pericytes microvasculatures after 4 days of treatment with TiO2 nanoparticles. The particles do not have a destabilising effect on networks formed. Red is F-actin staining and green is CD31. Scale bar: 100 µm. B) Tagged RNA-decorated cationic silica nanoparticles were injected after 10 days of vasculature maturation. Epifluorescence and bright field images were acquired at three time points (1hr, 24 and 48 hours). Particles can be found in both HUVECs and HUVECs/Pericytes vasculatures at 24 hrs. Scale bar: 100 µm.

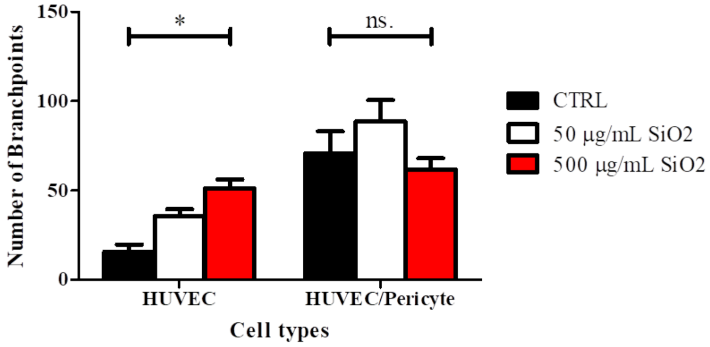


**Figure S11. Impact of PDMAEMA-coated $SiO_2$ nanoparticles on vascular network integrity.** Quantification of number of branchpoints in mono- and co-cultures after 4 days incubation with PDMAEMA-brush coated nanoparticles.

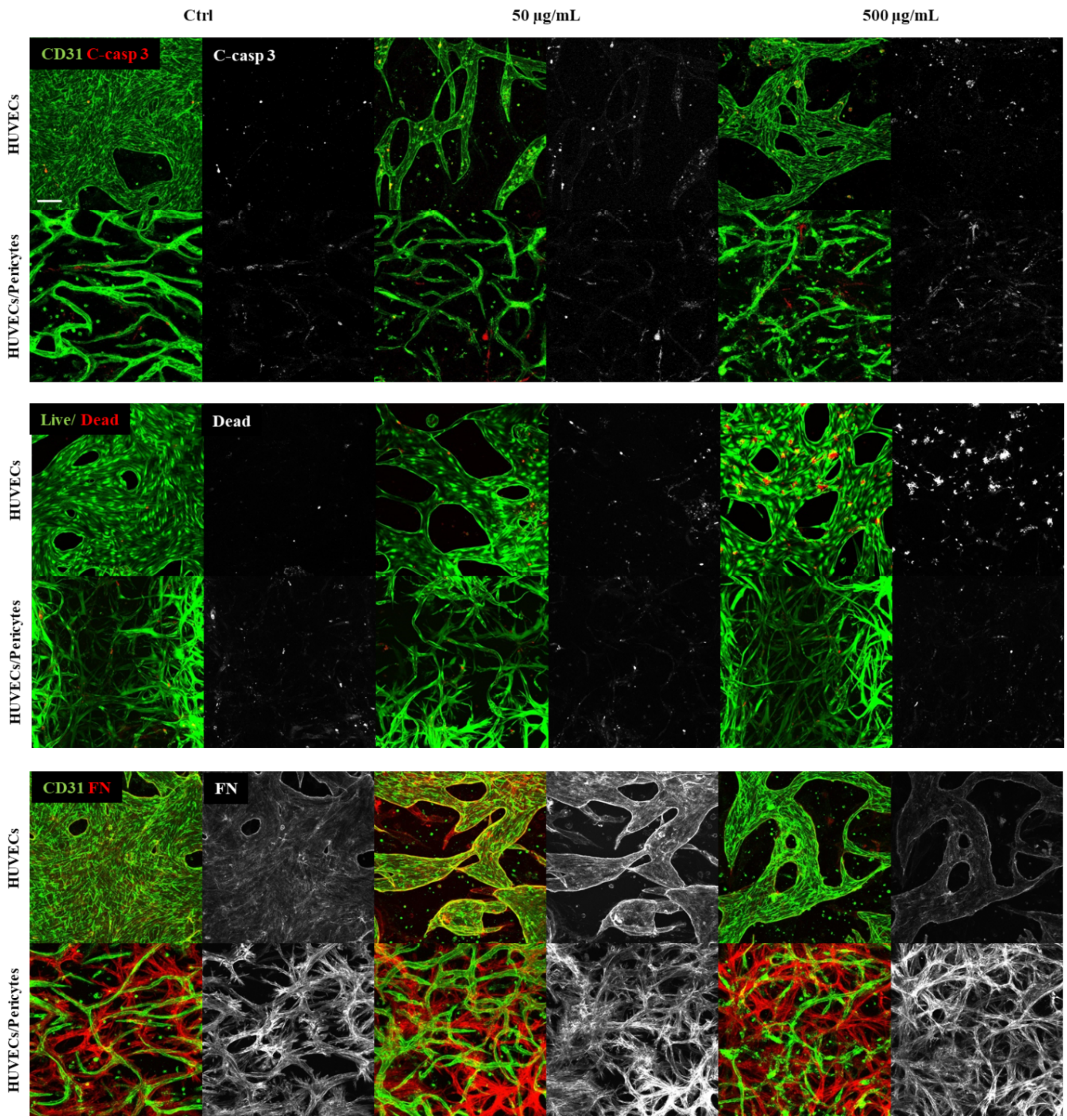


**Figure S12. Apoptosis, cytotoxicity and FN deposition after PDMAEMA-coated silica nanoparticles incubation.** Confocal images showing apoptosis (cleaved-caspase 3), cytotoxicity (live/dead assay) and FN deposition in mono and co- culture after treatment with 50 and 500 µg/mL cationic nanoparticles. Scale bar is 100 µm.
